# RNA sequencing (RNA-seq) reveals extremely low levels of reticulocyte-derived globin gene transcripts in peripheral blood from horses (*Equus caballus*) and cattle (*Bos taurus*)

**DOI:** 10.1101/294330

**Authors:** Carolina N. Correia, Kirsten E. McLoughlin, Nicolas C. Nalpas, David A. Magee, John A. Browne, Kevin Rue-Albrecht, Stephen V. Gordon, David E. MacHugh

## Abstract

RNA-seq has emerged as an important technology for measuring gene expression in peripheral blood samples collected from humans and other vertebrate species. In particular, transcriptomics analyses of whole blood can be used to study immunobiology and develop novel biomarkers of infectious disease. However, an obstacle to these methods in many mammalian species is the presence of reticulocyte-derived globin mRNAs in large quantities, which can complicate RNA-seq library sequencing and impede detection of other mRNA transcripts. A range of supplementary procedures for targeted depletion of globin transcripts have, therefore, been developed to alleviate this problem. Here, we use comparative analyses of RNA-seq data sets generated from human, porcine, equine and bovine peripheral blood to systematically assess the impact of globin mRNA on routine transcriptome profiling of whole blood in cattle and horses. The results of these analyses demonstrate that total RNA isolated from equine and bovine peripheral blood contains very low levels of globin mRNA transcripts, thereby negating the need for globin depletion and greatly simplifying blood-based transcriptomic studies in these two domestic species.

## 1. Introduction

It is increasingly recognised that new technological approaches are urgently required for infectious disease diagnosis, surveillance and management in burgeoning domestic animal populations as livestock production intensifies across the globe (Thornton, 2010; Nabarro and Wannous, 2014; Animal Task Force, 2016). In this regard, new strategies have emerged that leverage peripheral blood gene expression to study host immunobiology and to identify panels of RNA transcript biomarkers that can be used as specific biosignatures of infection by particular pathogens for both animal and human infectious disease (Ramilo and Mejias, 2009; Mejias and Ramilo, 2014; Chaussabel, 2015; Ko et al., 2015; Holcomb et al., 2017). For example, we and others have applied this approach to bovine tuberculosis (BTB) caused by infection with *Mycobacterium bovis* (Meade et al., 2007; Killick et al., 2011; Blanco et al., 2012; Churbanov and Milligan, 2012; McLoughlin et al., 2014; Cheng et al., 2015). It is also important to note that peripheral blood transcriptomics using technologies such as microarrays or RNA-sequencing (RNA-seq) can be used to monitor changes in the physiological status of domestic animals due to reproductive status, diet and nutrition or stress (O’Loughlin et al., 2012; Takahashi et al., 2012; Song et al., 2013; Kolli et al., 2014; Shen et al., 2014; de Greeff et al., 2016; Elgendy et al., 2016; Jegou et al., 2016).

During the last 15 years, a major hindrance to whole blood transcriptomics studies has emerged, which is the presence of large quantities of globin mRNA transcripts in peripheral blood from many mammalian species (Wu et al., 2003; Fan and Hegde, 2005; Liu et al., 2006). This is a consequence of abundant globin and globin mRNA transcripts in circulating reticulocytes, which in humans, may account for more than 95% of the total cellular mRNA content in these immature erythrocytes (Debey et al., 2004). Reticulocytes, in turn, account for 1-4% of the erythrocytes in healthy adult humans, which corresponds to between 5 × 10^7^ and 2 × 10^8^ cells per ml compared to 7 × 10^6^ cells per ml for leukocytes (Greer et al., 2013). Hence, globin transcripts can account for a substantial proportion of total detectable mRNAs in peripheral blood samples collected from humans and many other mammals (Bruder et al., 2010; Winn et al., 2010; Schwochow et al., 2012; Choi et al., 2014; Shin et al., 2014; Bowyer et al., 2015; Huang et al., 2016; Morey et al., 2016). In particular, for humans, more than 70% of peripheral blood mRNA transcripts are derived from the haemoglobin subunit alpha 1, subunit alpha 2 and subunit beta genes *(HBA1, HBA2* and *HBB)* (Wu et al., 2003; Field et al., 2007; Mastrokolias et al., 2012).

The emergence of massively parallel transcriptome profiling for clinical applications in human peripheral blood—initially with gene expression microarrays, but more recently using RNA-seq— has prompted development of methods for the systematic reduction of globin mRNAs in total RNA samples purified from peripheral blood samples, including: oligonucleotides that bind to globin mRNA molecules with subsequent digestion of the RNA strand of the RNA:DNA hybrid (Wu et al., 2003); peptide nucleic acid (PNA) oligonucleotides that are complementary to globin mRNAs and block reverse transcription of these targets (Liu et al., 2006); the GLOBINclear™ system, which uses biotinylated oligonucleotides that hybridise with globin transcripts followed by capture and separation using streptavidin-coated magnetic beads (Field et al., 2007); and the recently introduced GlobinLock method that uses a pair of modified oligonucleotides complementary to the 3’ portion of globin transcripts and that block enzymatic extension (Krjutskov et al., 2016).

In the present study we use RNA-seq data generated from globin-depleted and non-depleted total RNA purified from human and porcine peripheral blood, in conjunction with non-depleted total RNA isolated from equine and bovine peripheral blood, for a comparative investigation of the impact of reticulocyte-derived globin mRNA transcripts on routine transcriptome profiling of blood in domestic cattle and horses. The primary objective of the present study to test the hypothesis that both cattle and horses exhibit significantly lower quantities of haemoglobin gene transcripts compared to humans and pigs.

## 2. Materials and Methods

### 2.1. Data sources

RNA-seq data sets from human peripheral whole blood samples used for assessment of globin depletion and with parallel non-depleted controls (Shin et al., 2014) were obtained from the NCBI Gene Expression Omnibus (GEO) database (accession number GSE53655). A comparable RNA-seq data set from globin-depleted and non-depleted porcine peripheral whole blood was obtained directly from the study authors (Choi et al., 2014). A published RNA-seq data set (Ropka-Molik et al., 2017) from equine non-depleted peripheral whole blood was obtained from the NCBI GEO database (accession number GSE83404). Finally, bovine RNA-seq data from peripheral whole blood were generated by us as described below and can be obtained from the European Nucleotide Archive (ENA) database (accession number to be determined). A summary overview of the methodology used for the current study is shown in **Figure 1**.

**FIGURE 1.**
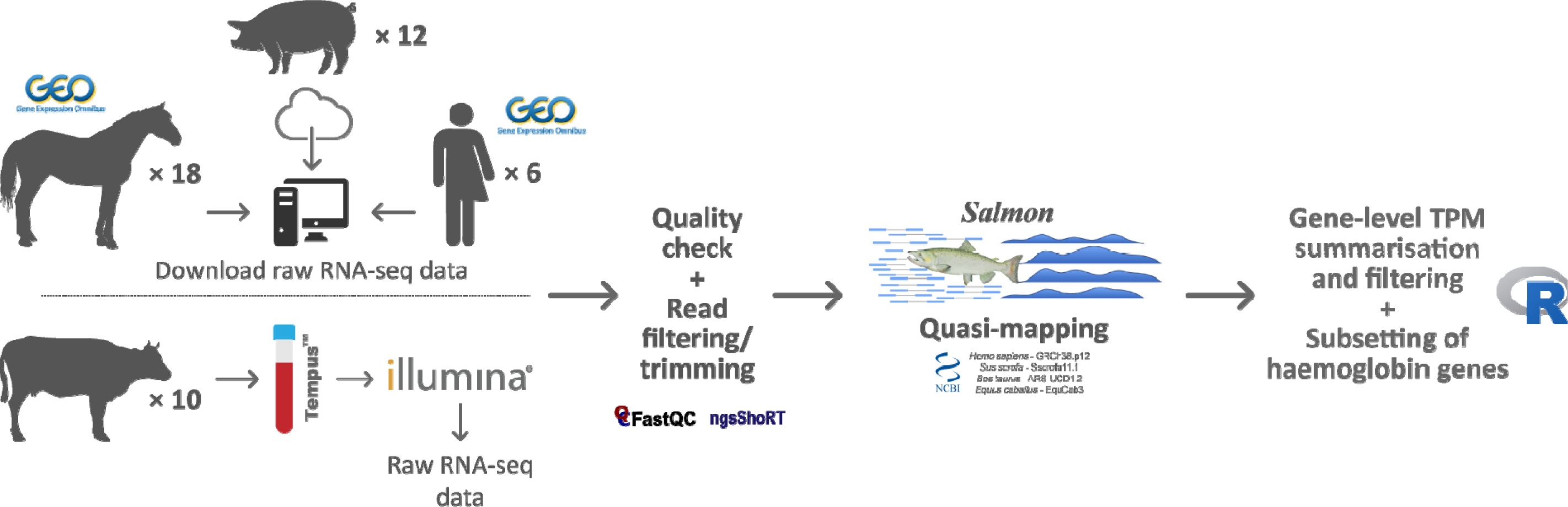
Schematic of the bioinformatics workflow for RNA-seq data acquisition, quality control, analysis and interpretation.

### 2.2. Human, porcine and equine sample collection, globin depletion and RNA-seq libraries

Detailed information concerning ethics approval, sample collection, total RNA extraction, and RNA-seq library preparation and sequencing for the human, porcine, and equine data sets is provided in the original publications (Choi et al., 2014; Shin et al., 2014; Ropka-Molik et al., 2017). Supplementary Table 1 provides summary information on the human, porcine and equine samples and RNA-seq libraries.

**Table 1:**
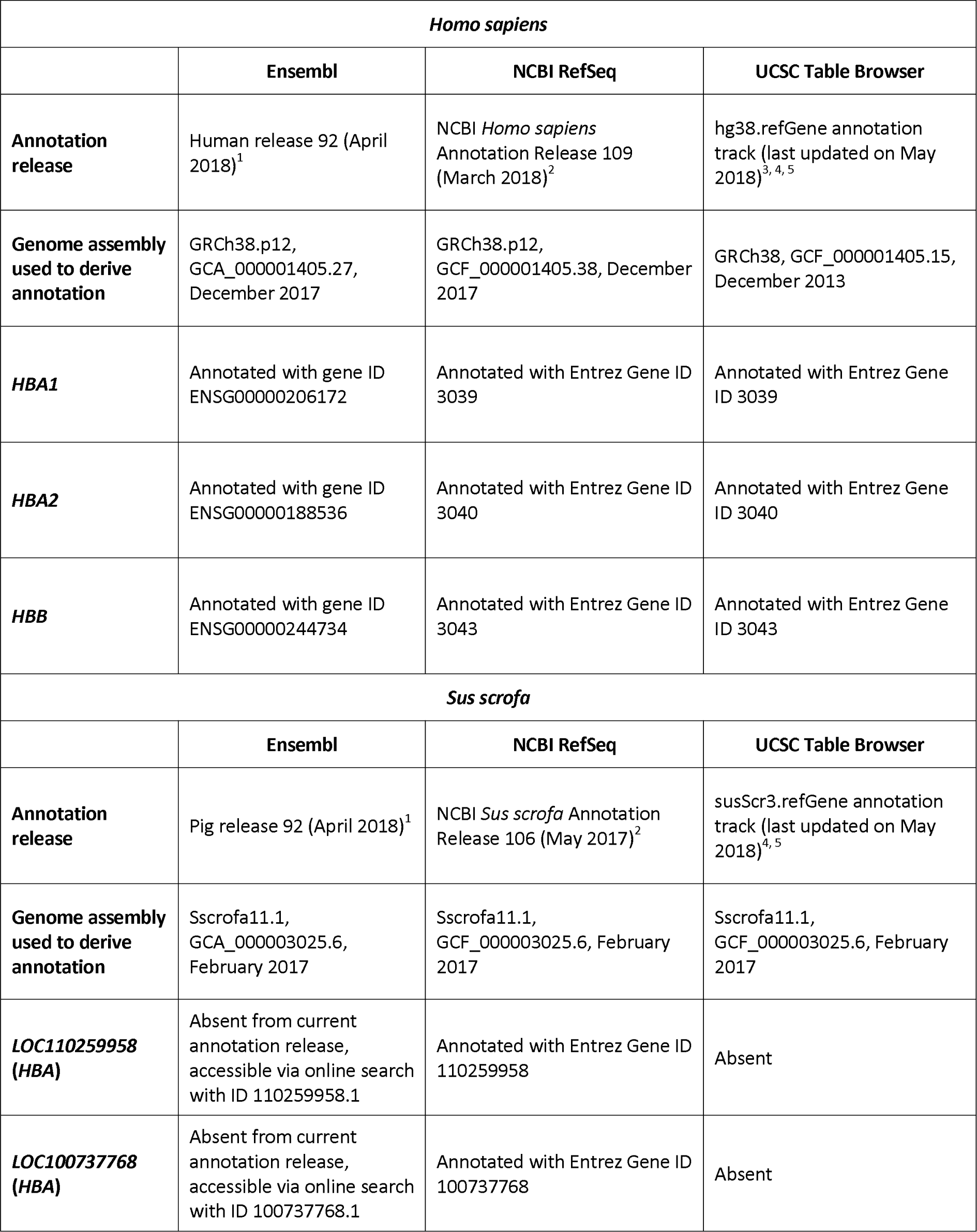
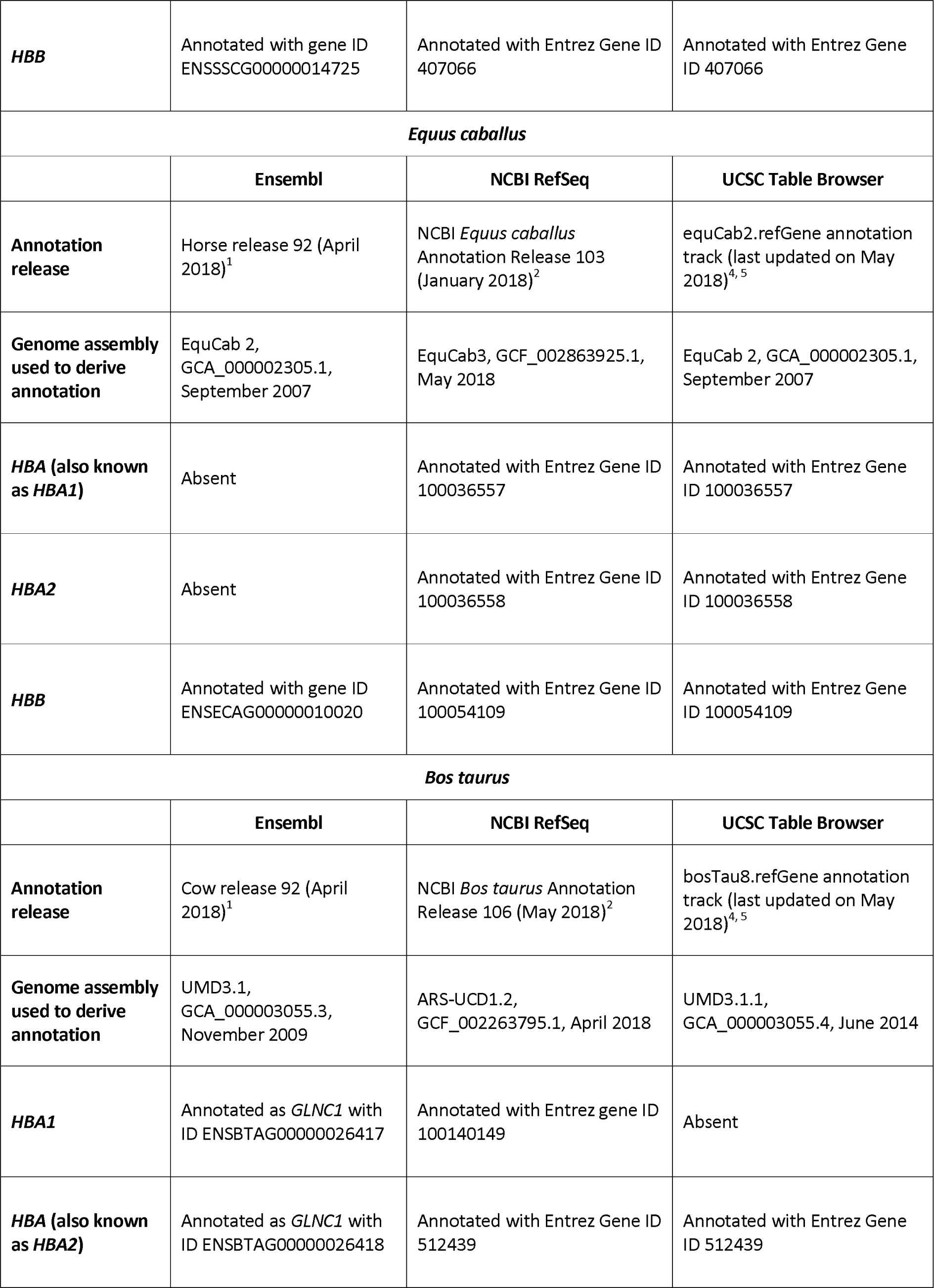
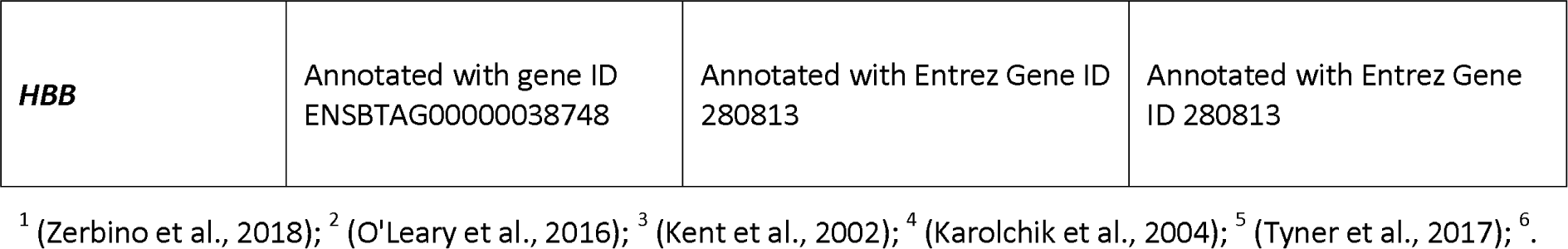
Status of current human, porcine, equine and bovine haemoglobin gene annotations in the Ensembl, NCBI RefSeq, and UCSC databases.

| *Homo sapiens* |  |  |  |
| --- | --- | --- | --- |
|  | Ensembl | NCBI RefSeq | UCSC Table Browser |
| **Annotation release** | Human release 92 (April 2018)<sup>1</sup> | NCBI *Homo sapiens* Annotation Release 109 (March 2018)<sup>2</sup> | hg38.refGene annotation track (last updated on May 2018)<sup>3, 4, 5</sup> |
| **Genome assembly used to derive annotation** | GRCh38.p12, GCA_000001405.27, December 2017 | GRCh38.p12, GCF_000001405.38, December 2017 | GRCh38, GCF_000001405.15, December 2013 |
| **HBA1** | Annotated with gene ID ENSG00000206172 | Annotated with Entrez Gene ID 3039 | Annotated with Entrez Gene ID 3039 |
| **HBA2** | Annotated with gene ID ENSG00000188536 | Annotated with Entrez Gene ID 3040 | Annotated with Entrez Gene ID 3040 |
| **HBB** | Annotated with gene ID ENSG00000244734 | Annotated with Entrez Gene ID 3043 | Annotated with Entrez Gene ID 3043 |
| *Sus scrofa* |  |  |  |
|  | Ensembl | NCBI RefSeq | UCSC Table Browser |
| **Annotation release** | Pig release 92 (April 2018)<sup>1</sup> | NCBI *Sus scrofa* Annotation Release 106 (May 2017)<sup>2</sup> | susScr3.refGene annotation track (last updated on May 2018)<sup>4, 5</sup> |
| **Genome assembly used to derive annotation** | Sscrofa11.1, GCA_000003025.6, February 2017 | Sscrofa11.1, GCF_000003025.6, February 2017 | Sscrofa11.1, GCF_000003025.6, February 2017 |
| **LOC110259958 (HBA)** | Absent from current annotation release, accessible via online search with ID 110259958.1 | Annotated with Entrez Gene ID 110259958 | Absent |
| **LOC100737768 (HBA)** | Absent from current annotation release, accessible via online search with ID 100737768.1 | Annotated with Entrez Gene ID 100737768 | Absent |
| **Impact of globin transcripts on RNA-seq transcriptomics using equine and bovine peripheral blood** |  |  |  |
| **HBB** | Annotated with gene ID ENSSCG00000014725 | Annotated with Entrez Gene ID 407066 | Annotated with Entrez Gene ID 407066 |
| ***Equus caballus*** |  |  |  |
|  | **Ensembl** | **NCBI RefSeq** | **UCSC Table Browser** |
| **Annotation release** | Horse release 92 (April 2018)<sup>1</sup> | NCBI *Equus caballus* Annotation Release 103 (January 2018)<sup>2</sup> | equCab2.refGene annotation track (last updated on May 2018)<sup>4,5</sup> |
| **Genome assembly used to derive annotation** | EquCab 2, GCA_000002305.1, September 2007 | EquCab3, GCF_002863925.1, May 2018 | EquCab 2, GCA_000002305.1, September 2007 |
| **HBA (also known as HBA1)** | Absent | Annotated with Entrez Gene ID 100036557 | Annotated with Entrez Gene ID 100036557 |
| **HBA2** | Absent | Annotated with Entrez Gene ID 100036558 | Annotated with Entrez Gene ID 100036558 |
| **HBB** | Annotated with gene ID ENSECAG00000010020 | Annotated with Entrez Gene ID 100054109 | Annotated with Entrez Gene ID 100054109 |
| ***Bos taurus*** |  |  |  |
|  | **Ensembl** | **NCBI RefSeq** | **UCSC Table Browser** |
| **Annotation release** | Cow release 92 (April 2018)<sup>1</sup> | NCBI *Bos taurus* Annotation Release 106 (May 2018)<sup>2</sup> | bosTau8.refGene annotation track (last updated on May 2018)<sup>4,5</sup> |
| **Genome assembly used to derive annotation** | UMD3.1, GCA_000003055.3, November 2009 | ARS-UCD1.2, GCF_002263795.1, April 2018 | UMD3.1.1, GCA_000003055.4, June 2014 |
| **HBA1** | Annotated as *GLNC1* with ID ENSBTAG00000026417 | Annotated with Entrez gene ID 100140149 | Absent |
| **HBA (also known as HBA2)** | Annotated as *GLNC1* with ID ENSBTAG00000026418 | Annotated with Entrez Gene ID 512439 | Annotated with Entrez Gene ID 512439 |
| ***HBB*** | Annotated with gene ID<br>ENSBTAG00000038748 | Annotated with Entrez Gene ID<br>280813 | Annotated with Entrez Gene<br>ID 280813 |

<sup>1</sup> (Zerbino et al., 2018); <sup>2</sup> (O'Leary et al., 2016); <sup>3</sup> (Kent et al., 2002); <sup>4</sup> (Karolchik et al., 2004); <sup>5</sup> (Tyner et al., 2017); <sup>6</sup>.

In brief, for the human samples, peripheral blood from six healthy subjects (three females and three males) was collected into PAXgene blood RNA tubes (PreAnalytiX/Qiagen Ltd., Manchester, UK). Total RNA, including small RNAs, was purified from the collected blood samples using the PAXgene Blood miRNA Kit (PreAnalytiX/Qiagen Ltd.) as described by Shin et al. (Shin et al., 2014). Human *HBA1, HBA2* and *HBB* mRNA transcripts were depleted from a subset of the total RNA samples using the GLOBINclear kit (Invitrogen™/Thermo Fisher Scientific, Loughborough, UK). RNA-seq data was then generated using 24 paired-end (PE) RNA-seq libraries (12 undepleted and 12 globin-depleted) generated from the six biological replicates and six identical technical replicates created from pooled total RNA across all six donor samples. The multiplexing and sequencing was then performed such that data for the 12 samples in each treatment group (undepleted and globin depleted) was generated from two separate lanes of a single flow cell twice, for a total of four sequencing lanes (Shin et al., 2014).

Porcine peripheral blood samples were collected from 12 healthy crossbred pigs (Duroc × (Landrace × Yorkshire)) using Tempus™ blood RNA tubes (Applied Biosystems™/Thermo Fisher Scientific, Warrington, UK) and total RNA was purified using the MagMAX™ for Stabilized Blood Tubes RNA Isolation Kit (Invitrogen™/Thermo Fisher Scientific) (Choi et al., 2014). Porcine *HBA* and *HBB* mRNA transcripts were subsequently depleted from a subset of the total RNA samples using a modified RNase H globin depletion method with custom porcine-specific antisense oligonucleotides for *HBA* and *HBB.* RNA-seq data was then generated from 24 PE RNA-seq libraries (12 undepleted and 12 globin-depleted).

Equine peripheral blood samples were collected using Tempus™ blood RNA tubes from 12 healthy Arabian horses (five females and seven males) at three different time points during flat racing training (Ropka-Molik et al., 2017). In addition, peripheral blood samples were collected from six healthy untrained Arabian horses (two females and four males). Total RNA was purified using the MagMAX™ for Stabilized Blood Tubes RNA Isolation Kit and 37 of the 42 total RNA samples were used to generate single-end (SE) libraries for RNA-seq data generation. Globin depletion for the equine samples was not performed prior to RNA-seq library preparation (Katarzyna Ropka-Molik, pers. comm.).

### 2.3. Bovine peripheral blood collection and RNA extraction

Approximately 3 ml of peripheral blood from ten age-matched healthy male Holstein-Friesian calves were collected into Tempus™ blood RNA tubes. The Tempus™ Spin RNA Isolation Kit (Applied Biosystems™/Thermo Fisher Scientific) was used to perform total RNA extraction and purification, following the manufacturer’s instructions. RNA quantity and quality checking were performed using a NanoDrop™ 1000 spectrophotometer (Thermo Fisher Scientific, Waltham, MA, USA) and an Agilent 2100 Bioanalyzer using an RNA 6000 Nano LabChip kit (Agilent Technologies Ltd., Cork, Ireland). The majority of samples displayed a 260/280 ratio greater than 1.8 and an RNA integrity number (RIN) greater than 8.0 (Supplementary Table 2). Globin mRNA depletion was not performed on the total RNA samples purified from bovine peripheral blood samples.

### 2.4. Bovine RNA-seq library generation and sequencing

Individually barcoded strand-specific RNA-seq libraries were prepared with 1 μg of total RNA from each sample. Two rounds of poly(A)^+^ RNA purification were performed for all RNA samples using the Dynabeads^®^ mRNA DIRECT™ Micro Kit (Thermo Fisher Scientific) according to the manufacturer’s instructions. The purified poly(A)^+^ RNA was then used to generate strand-specific RNA-seq libraries using the ScriptSeq™ v2 RNA-Seq Library Preparation Kit, the ScriptSeq™ Index PCR Primers (Sets 1 to 4) and the FailSafe™ PCR enzyme system (all sourced from Epicentre^®^/Illumina^®^ Inc., Madison, WI, USA), according to the manufacturer’s instructions.

RNA-seq libraries were purified using the Agencourt^®^ AMPure^®^ XP system (Beckman Coulter Genomics, Danvers, MA, USA) according to the manufacturer’s instructions for double size selection (0.75× followed by 1.0× ratio). RNA-seq libraries were quantified using a Qubit^®^ fluorometer and Qubit^®^ dsDNA HS Assay Kit (Invitrogen™/Thermo Fisher Scientific), while library quality checks were performed using an Agilent 2100 Bioanalyzer and High Sensitivity DNA Kit (Agilent Technologies Ltd.). Individually barcoded RNA-seq libraries were pooled in equimolar quantities and the quantity and quality of the final pooled libraries (three pools in total) were assessed as described above. Cluster generation and high-throughput sequencing of three pooled RNA-seq libraries were performed using an Illumina^®^ HiSeq™ 2000 Sequencing System at the MSU Research Technology Support Facility (RTSF) Genomics Core (https://rtsf.natsci.msu.edu/genomics; Michigan State University, MI, USA). Each of the three pooled libraries were sequenced independently on five lanes split across multiple Illumina^®^ flow cells. The pooled libraries were sequenced as PE 2 × 100 nucleotide reads using Illumina^®^ version 5.0 sequencing kits.

Deconvolution (filtering and segregation of sequence reads based on the unique RNA-seq library barcode index sequences; Supplementary Table 2) was performed by the MSU RTSF Genomics Core using a pipeline that simultaneously demultiplexed and converted pooled sequence reads into discrete FASTQ files for each RNA-seq sample with no barcode index mismatches permitted. The RNA-seq FASTQ sequence read data for the bovine samples were obtained from the MSU RTSF Genomics Core FTP server.

### 2.5. RNA-seq data quality control and filtering/trimming of reads

Bioinformatics procedures and analyses were performed as described below for the human, porcine, equine, and bovine samples, except were specifically indicated. All of the bioinformatics workflow scripts were developed using GNU bash (version 4.3.48) (Free Software Foundation, 2013), Python (version 3.5.2) (Python Software Foundation, 2017), and R (version 3.4.0) (R Core Team, 2017). The scripts and further information are available at a public GitHub repository (https://github.com/carolcorreia/GlobinRNA-sequencing). Computational analyses were performed on a 32-core Linux Compute Server (4× AMD Opteron™ 6220 processors at 3.0 GHz with 8 cores each), with 256 GB of RAM, 24 TB of hard disk drive storage, and with Ubuntu Linux OS (version 14.04.4 LTS). Deconvoluted FASTQ files (generated from SE equine RNA-seq libraries and PE RNA-seq libraries for the other species) were quality-checked with FastQC (version 0.11.5) (Andrews, 2016).

Using the ngsShoRT software package (version 2.2) (Chen et al., 2014), filtering/trimming consisted of: (1) removal of SE or PE reads with adapter sequences (with up to three mismatches); (2) removal of SE or PE reads of poor quality (i.e., at least one of the reads containing ≥ 25% bases with a Phred quality score below 20); (3) for porcine samples only, 10 bases were trimmed at the 3’ end of all reads; (4) removal of SE or PE reads that did not meet the required minimum length (70 nucleotides for human and equine, 80 nucleotides for porcine and 100 nucleotides for bovine). Filtered/trimmed FASTQ files were then re-evaluated using FastQC. Filtered FASTQ files were transferred to a 36-core/64-thread Compute Server (2× Intel^®^ Xeon^®^ CPU E5-2697 v4 at 2.30 GHz with 18 cores each), with 512 GB of RAM, 96 TB SAS storage (12× 8 TB at 7200 rpm), 480 GB SSD storage, and with Ubuntu Linux OS (version 16.04.2 LTS).

### 2.6. Transcript quantification

The Salmon software package (version 0.8.2) (Patro et al., 2017) was used in quasi-mapping-mode for transcript quantification. Sequence-specific and fragment-level GC bias correction was enabled and transcript abundance was quantified in transcripts per million (TPM) for each filtered library (multiple lanes from the same library were processed together) was estimated after mapping of SE or PE reads to their respective reference transcriptomes. As summarised in **Table 1**, the NCBI RefSeq database is currently the only one to contain haemoglobin gene annotations for all species analysed. Hence, NCBI RefSeq reference transcript models were used for the human, porcine, equine, and bovine data sets. Detailed information about these reference transcriptomes is provided in Supplementary Table 3.

### 2.7. Gene annotations and summarisation of TPM estimates at the gene level

Using R (3.5.0) within the RStudio IDE (version 1.1.447) (RStudio Team, 2015) and Bioconductor (version 3.7 using BiocInstaller 1.30.0) (Gentleman et al., 2004), the GenomicFeatures (version 1.32.0) (Lawrence et al., 2013) and AnnotationDbi (version 1.42.1) (Pagès et al., 2017) packages were used to obtain corresponding gene and transcript identifiers from the NCBI RefSeq annotation releases pertinent to each species, as detailed in **Table 1**. Using these identifiers, the tximport (version 1.8.0) package (Soneson et al., 2015) was used to import into R and summarise at gene level the TPM estimates obtained from the Salmon tool. A threshold of greater than or equal to 1 TPM across at least half of the total number of samples (≥ 12 for human and porcine, ≥18 for equine, and ≥5 for bovine) was applied in order to remove lowly expressed genes.

### 2.8. Data exploration, plotting and summary statistics

Data wrangling and tidying from all species was performed using the following R packages: tidyverse (version 1.2.1) (Wickham, 2017b), dplyr (version 0.7.5) (Wickham et al., 2017), tidyr (version 0.8.1) (Wickham and Henry, 2017), reshape2 (version 1.4.3) (Wickham, 2017a), and magrittr (version 1.5) (Bache and Wickham, 2017). The ggplot2 (version 2.2.1) (Wickham and Chang, 2017), and ggjoy (version 0.4.1) (Wilke, 2017), packages were used for figure generation. Finally, the mean and standard deviation were calculated for the undepleted and globin-depleted groups in each species using the skimr (version 1.0.2) R package (McNamara et al., 2017).

## 3. Results and Discussion

### 3.1. Status of human, porcine, equine and bovine haemoglobin gene annotations

Annotation of the haemoglobin subunit alpha 1 and 2 genes *(HBA1* and *HBA2*, respectively) is well established for the human genome; however, annotations for these genes in the porcine, equine and bovine genomes are inconsistent across databases. As shown in **Table 1**, the porcine *HBA* gene annotation is absent from Ensembl and the UCSC Table Browser. For the NCBI RefSeq database, this gene has been assigned to two loci *(LOC110259958* and *LOC100737768)* that have similar descriptions (haemoglobin subunit alpha and haemoglobin subunit alpha-like). Therefore, these NCBI LOC symbols were used.

Equine *HBA (HBA1)* and *HBA2* genes are absent from the current Ensembl annotation release. Similarly, bovine *HBA1* and *HBA (HBA2)* have been annotated as *GLNC1* in Ensembl, whereas *HBA1* is absent from the UCSC Table Browser annotation (**Table 1**). In the NCBI RefSeq database, equine *HBA (HBA1)* is described as haemoglobin subunit alpha 1; and bovine *HBA (HBA2)* is described as haemoglobin subunit alpha 2, thus their descriptions are shown in parenthesis herein. In contrast to these observations, haemoglobin subunit beta *(HBB)* genes for the four species are well annotated in Ensembl, NCBI RefSeq and UCSC Genome Browser databases (**Table 1**).

At the time of writing, NCBI RefSeq is the only database that contains annotations for all three haemoglobin genes in all species analysed. Additionally, equine and bovine gene annotations are based on the latest genome assemblies (**Table 1**). EquCab3 and ARS-UCD1.2 have incorporated major improvements compared to previous versions, including increased genome coverage (from 6.8× and 9×, to 80×, respectively), and incorporation of PacBio sequencing reads (Kalbfleisch et al., 2018; Rosen et al., 2018).

### 3.2. Basic RNA-seq data outputs

Unfiltered SE (equine libraries) or PE (human, porcine, and bovine libraries) RNA-seq FASTQ files were quality-checked, adapter- and quality-filtered prior to transcript quantification. As shown in **Table 2**, the human and porcine undepleted groups each had approximately 40 million (M) raw reads per library, whereas globin-depleted libraries showed a mean of approximately 37 M and 31 M, respectively. Equine and bovine libraries, which did not include a globin depletion step had an average of 24 M raw reads and 21 M raw read pairs, respectively.

**Table 2:** Summary of RNA-seq filtering/trimming and mapping statistics.

| Species | Treatment | RNA-seq library type | Sequencing mode | Mean no. of reads (SE) or pairs (PE) | Mean no. of reads (SE) or pairs (PE) removed | Mean proportion of reads (SE) or pairs (PE) removed | Mean no. of observed fragments* | Mean no. of mapped fragments* | Average mapping rate | Reference source |
| --- | --- | --- | --- | --- | --- | --- | --- | --- | --- | --- |
| *Homo sapiens* | Undepleted | Inward unstranded | PE | 40,218,886 | 8,203,217 | 20.4% | 32,015,669 | 25,593,239 | 80.9% | NCBI RefSeq |
| *Homo sapiens* | Globin depleted | Inward unstranded | PE | 36,874,759 | 10,704,088 | 29.0% | 26,170,671 | 20,371,624 | 75.8% | NCBI RefSeq |
| *Sus scrofa* | Undepleted | Inward unstranded | PE | 39,036,515 | 4,613,991 | 11.8% | 28,685,437 | 25,129,140 | 87.7% | NCBI RefSeq |
| *Sus scrofa* | Globin depleted | Inward unstranded | PE | 31,339,886 | 3,899,427 | 12.4% | 22,867,049 | 19,959,995 | 87.3% | NCBI RefSeq |
| *Equus caballus* | Undepleted | Unstranded | SE | 24,271,141 | 38,892 | 0.2% | 14,850,797 | 11,387,774 | 76.5% | NCBI RefSeq |
| *Bos taurus* | Undepleted | Inward stranded forward | PE | 20,495,983 | 3,474,597 | 17.0% | 17,021,386 | 12,353,147 | 72.6% | NCBI RefSeq |

\*The Salmon tool categorises fragments as single read (for SE RNA-seq libraries) or a read pair (for PE RNA-seq libraries).

After adapter- and quality-filtering of RNA-seq libraries, an average of 20% and 29% read pairs were removed from the human undepleted and globin-depleted libraries, respectively. Conversely, approximately 12% of read pairs were removed from each of the porcine undepleted and globin-depleted libraries. For the undepleted equine and bovine RNA-seq libraries, an average of 0.2% reads and 17% read pairs were removed, respectively. Detailed information on filtering/trimming of RNA-seq libraries from all species, including technical replicates from libraries sequenced over multiple lanes, is presented in Supplementary Table 4. All data sets exhibited a mean mapping rate greater than 70% (**Table 2**). Supplementary Tables 5 contain sample-specific RNA-seq mapping statistics.

### 3.3. Transcript quantification

Transcript-level TPM estimates generated using the Salmon tool were imported into the R environment and summarised at gene level with the package tximport (Soneson et al., 2015). Gene-level TPM estimates represent the sum of corresponding transcript-level TPMs and provide results that are more accurate and comprehensible than transcript-level estimates (Soneson et al., 2015). In the current study, gene-level TPM estimates are referred as TPM.

Filtering of lowly expressed genes (see **Section 2.7**) resulted in 12,951 genes expressed across all human samples, and represented 24% of 54,644 total annotated genes and pseudogenes. Porcine samples showed a total of 9,396 expressed genes (31% of 30,334 annotated genes and pseudogenes); and equine and bovine samples exhibited 12,724 (38% of 33,146) and 14,044 (40% of 35,143) expressed genes, respectively.

The density distribution of TPM values for the human and porcine samples improved after globin depletion; this is evident by the shift of gene detection levels towards greater log10 TPM values for the globin-depleted samples in **Figure 2**. In this regard, it is noteworthy that the undepleted bovine and equine samples also exhibited similar TPM density distributions to the human and porcine globin-depleted samples.

**FIGURE 2.**
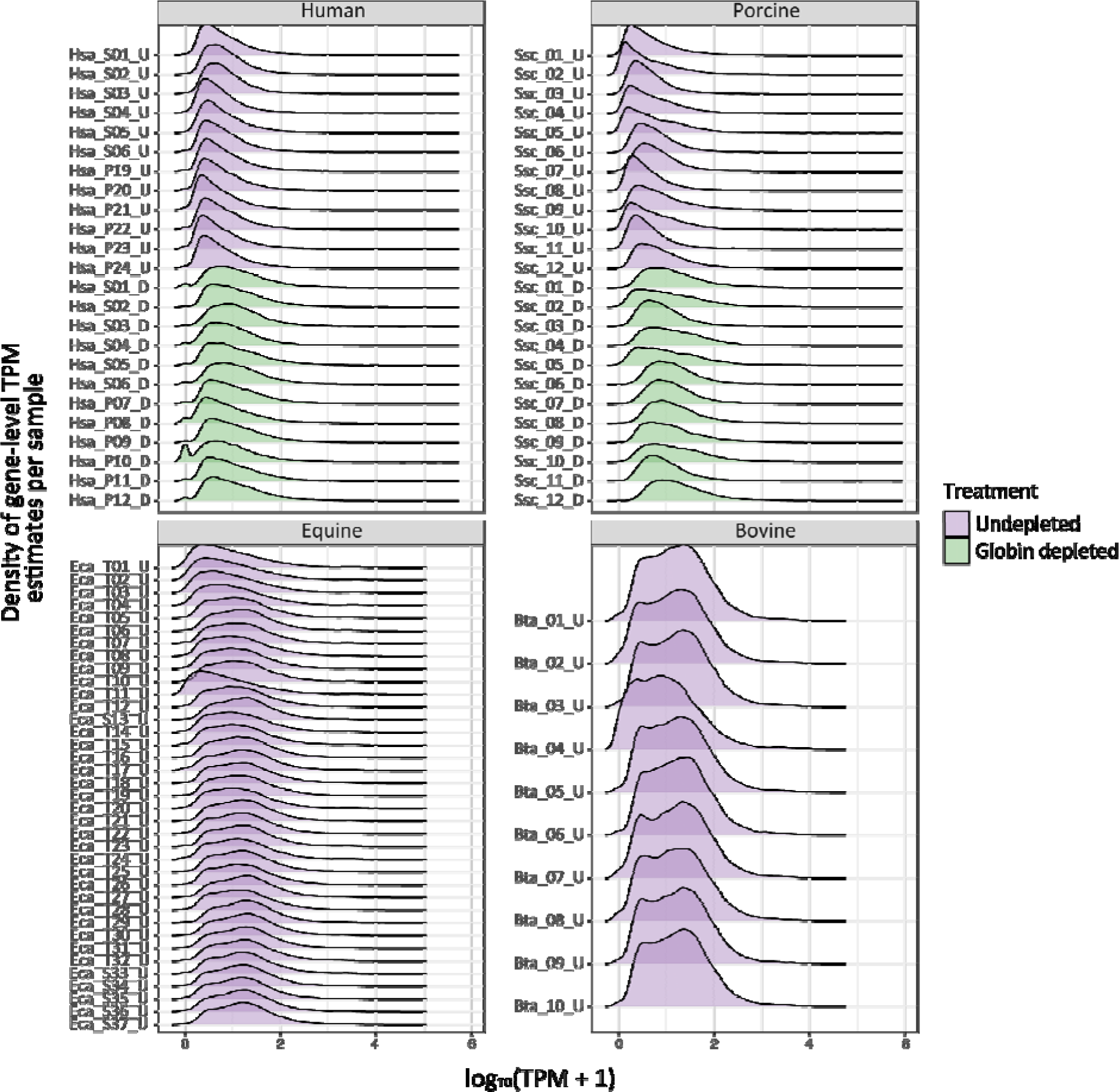
Ridge plots showing density of sample gene-level transcripts per million (TPM). Results are shown from undepleted (purple) or globin-depleted (green) treatments.

### 3.4. Proportions of human and porcine haemoglobin gene transcripts in undepleted and depleted peripheral blood

In line with previous reports (Field et al., 2007; Mastrokolias et al., 2012), the proportion of haemoglobin gene transcripts *(HBA1, HBA2*, and *HBB)* detected in undepleted human peripheral blood samples for the current study averaged 70% (**Figure 3** and Supplementary Table 6), which is lower than the mean proportion of 81% reported by Shin et al. (2014). On the other hand, after depletion the human samples exhibited an identical reduction to a 17% proportion of globin sequence reads in both the present study and that of Shin et al. (2014) (**Figure 3** and Supplementary Table 6).

**FIGURE 3.**
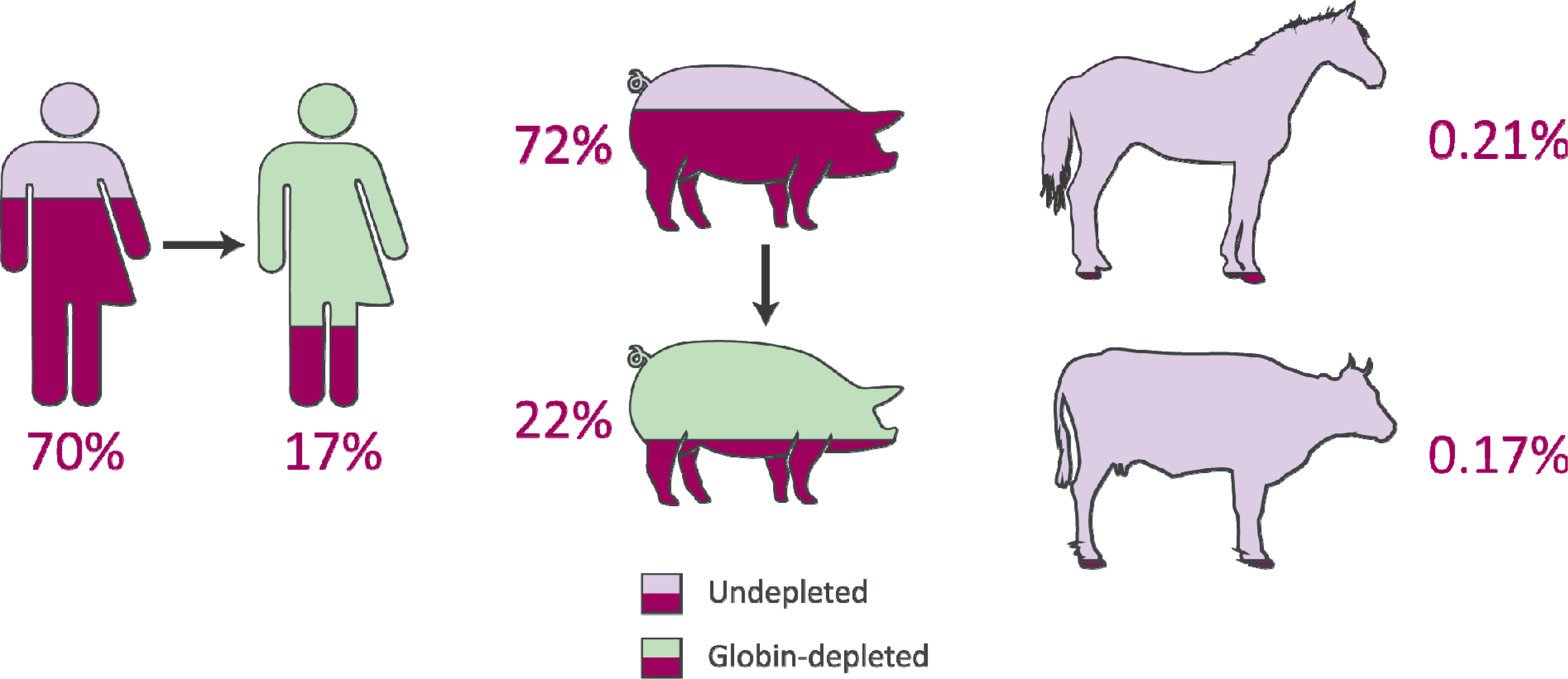
Average proportions of haemoglobin genes to total expressed genes from peripheral blood RNA-seq data in humans, pigs, horses and cattle.

In the current study, for the undepleted porcine peripheral blood samples, the percentage of haemoglobin gene transcripts *(LOC110259958* [HBA], *LOC100737768* [HBA], and *HBB)* observed as a proportion of the total expressed genes was 72% (**Figure 3** and Supplementary Table 6), which is considerably larger than the mean of 46.1% reported in the original study (Choi et al., 2014). Similarly, after depletion, the porcine samples in the present study contained a mean proportion of 22% globin transcripts (**Figure 3** and Supplementary Table 6) compared to a mean proportion of 8.9% reported by Choi et al. (2014). Additionally, **Table 3** shows the mean TPM for each haemoglobin gene across undepleted or globin-depleted samples.

**Table 3:** Summary statistics for haemoglobin gene-level transcripts per million (TPM).

| Species | Gene symbol | Treatment | No. of samples | Mean TPM | Standard deviation |
| --- | --- | --- | --- | --- | --- |
| *Homo sapiens* | *HBA1* | Undepleted | 12 | 191,209 | 16,601 |
| *Homo sapiens* | *HBA1* | Globin depleted | 12 | 66,718 | 23,557 |
| *Homo sapiens* | *HBA2* | Undepleted | 12 | 300,000 | 29,523 |
| *Homo sapiens* | *HBA2* | Globin depleted | 12 | 79,818 | 31,259 |
| *Homo sapiens* | *HBB* | Undepleted | 12 | 200,000 | 43,262 |
| *Homo sapiens* | *HBB* | Globin depleted | 12 | 20,770 | 6,706 |
| *Sus scrofa* | *LOC110259958 (HBA)* | Undepleted | 12 | 86 | 30 |
| *Sus scrofa* | *LOC110259958 (HBA)* | Globin depleted | 12 | 13 | 13 |
| *Sus scrofa* | *LOC100737768 (HBA)* | Undepleted | 12 | 243,864 | 31,605 |
| *Sus scrofa* | *LOC100737768 (HBA)* | Globin depleted | 12 | 84,021 | 86,095 |
| *Sus scrofa* | *HBB* | Undepleted | 12 | 476,284 | 52,939 |
| *Sus scrofa* | *HBB* | Globin depleted | 12 | 136,172 | 128,232 |
| *Equus caballus* | *HBA (HBA1)* | Undepleted | 37 | 443 | 560 |
| *Equus caballus* | *HBA2* | Undepleted | 37 | 653 | 789 |
| *Equus caballus* | *HBB* | Undepleted | 37 | 1,024 | 1,144 |
| *Bos taurus* | *HBA1* | Undepleted | 10 | 21 | 29 |
| *Bos taurus* | *HBA (HBA2)* | Undepleted | 10 | 1,101 | 1,102 |
| *Bos taurus* | *HBB* | Undepleted | 10 | 532 | 469 |

A number of possible explanations, including the different approaches used for read mapping and transcript quantification, may account for the different proportions of haemoglobin gene transcript detected in human and porcine samples for the present study compared to the original studies (Choi et al., 2014; Shin et al., 2014). For the present study, a recently developed lightweight alignment method was adopted (Salmon and tximport), in contrast to the more traditional methodologies used in the original publications. Shin and colleagues (2014) used the TopHat and Cufflinks software tools (Trapnell et al., 2012), while Choi et al. (2014) implemented TopHat with Htseq-count (Anders et al., 2015). In addition to this, different gene annotations were used: NCBI *Homo sapiens* Annotation Release 109 and NCBI *Sus scrofa* Annotation Release 106 were used for the present study, while UCSC hg18 *(Homo sapiens)* and Ensembl release 71 *(Sus scrofa)* were used by Shin et al. (2014) and Choi et al. (2014), respectively.

### 3.5. Equine and bovine peripheral blood contains extremely low levels of haemoglobin gene transcripts

The equine and bovine peripheral blood samples, which did not undergo globin depletion, had extremely low proportions of haemoglobin gene transcripts to total expressed genes: 0.21% and 0.17%, respectively (**Figure 3** and Supplementary Table 6). Notably, similar results have been reported in a transcriptomics study of bovine peripheral blood in response to vaccination against neonatal pancytopenia. In that study, 12 cows were profiled before and after vaccination (24 peripheral blood samples in total), and a mean proportion of 1.0% of RNA-seq reads were observed to map to the bovine α haemoglobin gene cluster on BTA25 or to the β haemoglobin gene cluster on BTA15 (Demasius et al., 2013). To the best of our knowledge, this is the first time that the average number of equine haemoglobin transcripts have been reported for RNA-seq data.

Finally, it is important to note that log2 TPM values for haemoglobin gene transcripts in the undepleted equine and bovine peripheral blood RNA samples are substantially lower than log_2_ TPM values for the globin-depleted human and porcine peripheral blood RNA samples (**Figure 4**). This is a direct consequence of extremely low levels of circulating reticulocytes in equine and bovine peripheral blood (Tablin and Weiss, 1985; Harper et al., 1994; Hossain et al., 2003; Cooper et al., 2005).

**FIGURE 4.**
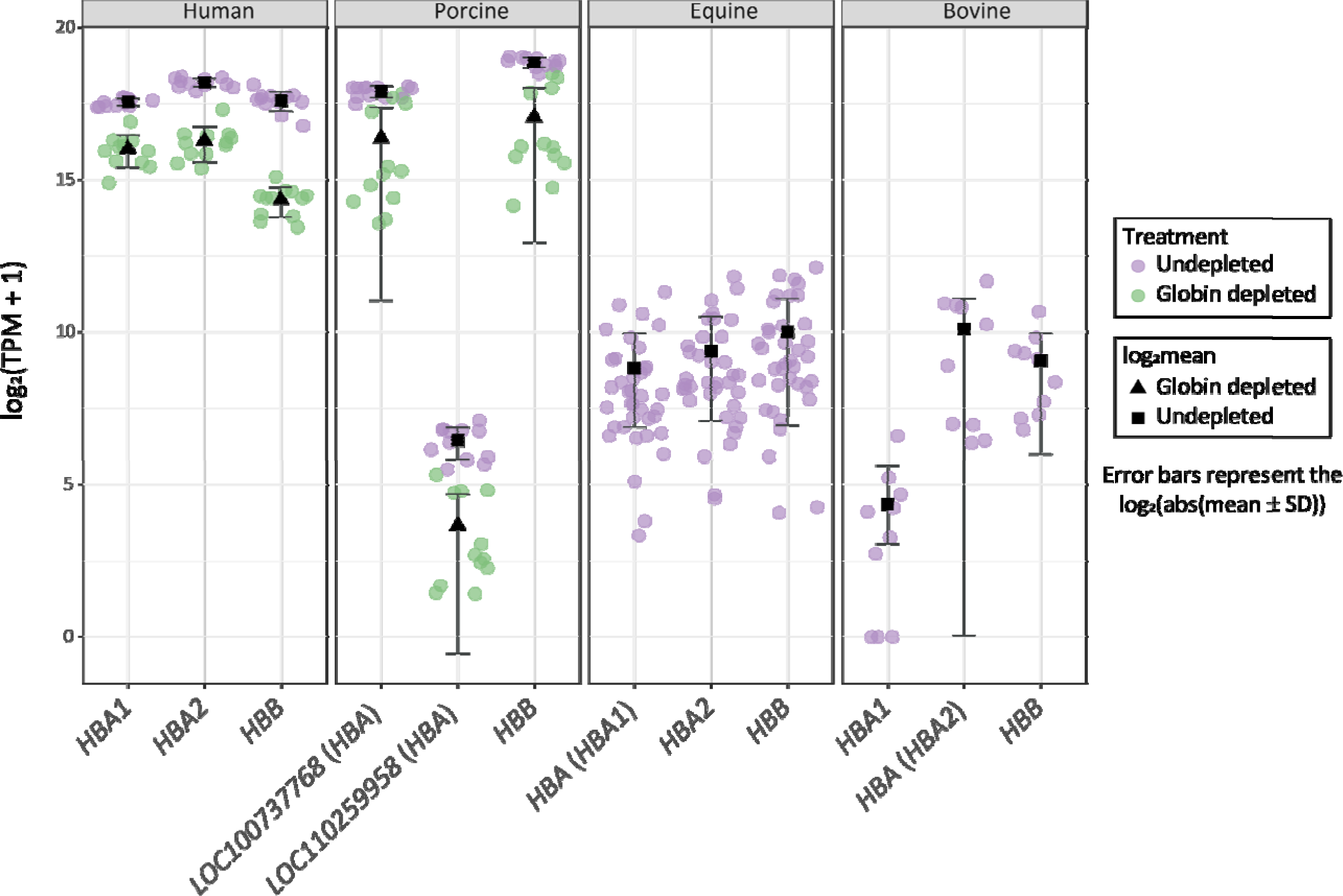
Distribution of haemoglobin gene-level transcripts per million (TPM). Results are shown from undepleted (purple) or globin-depleted (green) treatments.

### 3.6. Conclusion

In light of our RNA-seq data analyses, we propose that globin mRNA transcript depletion is not a pre-requisite for transcriptome profiling of bovine and equine peripheral blood samples. This observation greatly simplifies the laboratory and bioinformatics workflows required for RNA-seq studies of whole blood collected from domestic cattle and horses. It will also be directly relevant to future work on blood-based biomarker and biosignature development in the context of infectious disease, reproduction, nutrition and animal welfare. For example, transcriptomics of peripheral blood has been used extensively in development of new diagnostic and prognostic modalities for human tuberculosis (HTB) disease caused by infection with *Mycobacterium tuberculosis* (for reviews see: Blankley et al., 2014; Haas et al., 2016; Weiner and Kaufmann, 2017; Goletti et al., 2018). Therefore, as a consequence of this HTB research, comparable transcriptomics studies in cattle (Meade et al., 2007; Killick et al., 2011; Blanco et al., 2012; Churbanov and Milligan, 2012; McLoughlin et al., 2014; Cheng et al., 2015), and the ease with which RNA-seq can be performed in bovine peripheral blood, it should be feasible to develop transcriptomics-based biomarkers and biosignatures for bovine tuberculosis caused by *M. bovis* infection.

## 4. Data Accessibility

The RNA-seq data generated for this study using peripheral blood from ten age-matched healthy male Holstein-Friesian calves can be obtained from the ENA database (accession number to be determined).

## 6. Ethics Statement

Animal experimental work for the present study (cattle samples) was carried out according to the UK Animal (Scientific Procedures) Act 1986. The study protocol was approved by the Animal Health and Veterinary Laboratories Agency (AHVLA-Weybridge, UK), now the Animal & Plant Health Agency (APHA), Animal Use Ethics Committee (UK Home Office PCD number 70/6905).

## 7. Author Contributions

DEM, SVG, CNC, and KEM conceived and designed the project and organised bovine sample collection; KEM, NCN, DAM, and JAB performed RNA extraction and RNA-seq library generation; CNC, KEM, NCN, KRA, and DEM performed the analyses; CNC and DEM wrote the manuscript and all authors reviewed and approved the final manuscript.

## 9. Acknowledgements

The authors wish to express their gratitude to Prof Martin Vordermeier and Dr Bernardo Villarreal-Ramos (Animal and Plant Health Agency, UK) for provision of bovine peripheral blood samples, Prof Graham Plastow (University of Alberta, Canada) for provision of porcine peripheral blood RNA-seq data. We also thank Drs Gabriella Farries (University College Dublin) and Kerri Malone (EMBL-EBI, Cambridge, UK) for stimulating discussion and advice concerning genome annotations, equine genetics and data visualisation.

